# Seed variation impacts clustering stability in Single-Cell RNA-Seq and can be mitigated by StAbility-BasEd-Reassignment (SABER)

**DOI:** 10.64898/2026.06.16.732539

**Authors:** Nunzia Zagaria, Giulia Tini, Emanuele Bonetti, Luca Mazzarella

**Affiliations:** Department of Experimental Oncology, IEO, European Institute of Oncology IRCCS, Milan, Italy; Division of Advanced Molecular Diagnostics (DIMA), Department of Experimental Oncology, IEO, European Institute of Oncology IRCCS, Milan, Italy

## Abstract

Single-cell RNA-seq clustering is commonly treated as reproducible once a random seed is fixed, yet the choice of seed itself may alter cell assignments and downstream interpretation. We systematically quantified seed-induced clustering variability by running Louvain and Leiden clustering across 100 seeds in Seurat and Scanpy on 28 single-cell RNA-seq datasets from the Human Cell Atlas and IMMUcan. Using Element-Centric Consistency, we found that seed choice affected a substantial fraction of cells, with Scanpy showing more unstable assignments than Seurat on average, 40.46% versus 26.78% unstable cells, respectively. This increased stability came at a marked computational cost: Seurat required approximately 19-fold higher median memory than Scanpy. Seed-dependent clustering variability also propagated to cell-type annotation, particularly among transcriptionally related populations including macrophage/monocyte, endothelial/epithelial and T/NK cell states. To mitigate this instability, we developed StAbility-BasEd Reassignment (SABER), a Scanpy-based framework that identifies seed-sensitive cells across repeated clusterings and reassigns them to stable cluster cores using cosine similarity. SABER improved clustering quality while preserving annotation concordance and reduced median memory usage 3.5-fold compared with Seurat-Louvain. Our results identify seed choice as an underappreciated source of variability in single-cell analysis and provide a scalable strategy to improve clustering robustness.

## Introduction

In computational biology and machine learning, stochastic algorithms are often initialized with a fixed seed, a specific value that set random number generation algorithms. The practice of fixing a seed value is commonly used to support computational reproducibility, as it allows analyses to be rerun under the same initialization conditions ^1,2^. The pseudorandom numbers derived from the selected seed can influence different steps of an analysis, including parameters initialization, instance selection, and optimization trajectories. As a result, chaining only the seed can lead to divergent outcomes^3,4^. This “seed effect” has been the subject of investigation in a number of studies. Henderson and colleagues demonstrated that the use of different seeds across ten runs can result in the generation of statistically distinct distributions of algorithm performance^5^. Similar findings have been observed across a range of ML algorithms^6^, deep neural models^7^, and generative AI models for images^8^. Evaluating model robustness therefore requires testing multiple seeds to ensure results are not dependent on a particular initialization.

Single-cell RNA sequencing (scRNA-seq) pipelines^9^ involve several stochastic steps, particularly clustering, which identifies groups of transcriptionally similar cells. The most employed community detection methods are Louvain^10^ and Leiden^11^ algorithms, both of which identify communities by optimizing the modularity, which measures edge density within versus between communities. Leiden algorithm was developed as an improvement over Louvain by adding a refinement step that updates modularity exclusively for affected neighbors rather than all cells in every iteration, resulting in communities with enhanced density and superior computational efficiency in comparison to the Louvain algorithm. In both algorithms, the seed affects the node evaluation order during the initial modularity computation. Louvain and Leiden algorithms are implemented in widely adopted scRNA-seq tools: the R package Seurat^12^ and the Python library Scanpy^13^. These two packages show important differences in computational performance, that reflect the different memory allocation strategies adopted by their language environment.

Here, using publicly available datasets in the Human Cell Atlas (HCA)^14^ and IMMUcan^15^, we quantify the impact of seed choice across graph-based algorithms (Leiden vs Louvain) and commonly used analytical frameworks (Seurat vs Scanpy). We show that seed choice leads to highly variable degrees of cluster instability across datasets and analytical platforms, which generate potentially relevant inconsistencies. We developed StAbility-BasEd-Reassignment (SABER), a method that identifies cells whose cluster assignment is most sensitive to seed variation and reassigns them to stable clusters based on cosine similarity. This approach improves clustering stability while preserving biological structure and maintaining computational efficiency.

## Results

### Seed variation leads to clustering instability

To quantify seed variation in scRNA-seq clustering, we analyzed publicly available datasets from Human Cell Atlas (HCA)^14^ and IMMUcan^15^ databases (Supplementary Table 1). We applied Louvain and Leiden clustering in both Seurat and Scanpy across 100 distinct seeds (1-100). Three datasets could not be processed with Seurat for excessive memory usage despite using 190 GB of RAM and an additional 60 GB of swap memory, underscoring the substantial computational requirements of this framework. The remaining 28 datasets were retained for the final benchmark (Fig.1a). We quantified cluster stability using Element-Centric Consistency (ECC)^16^, a metric that measures how consistently individual cells are assigned to clusters across partitions. Cells with ECC ≤ 0.85 were defined as unstable. This threshold was experimentally derived from the IMMUcan datasets by identifying the median ECC value above which annotation became unstable in at least 5% of the seed iterations (Supplementary Fig. 1).

**Figure 1.**
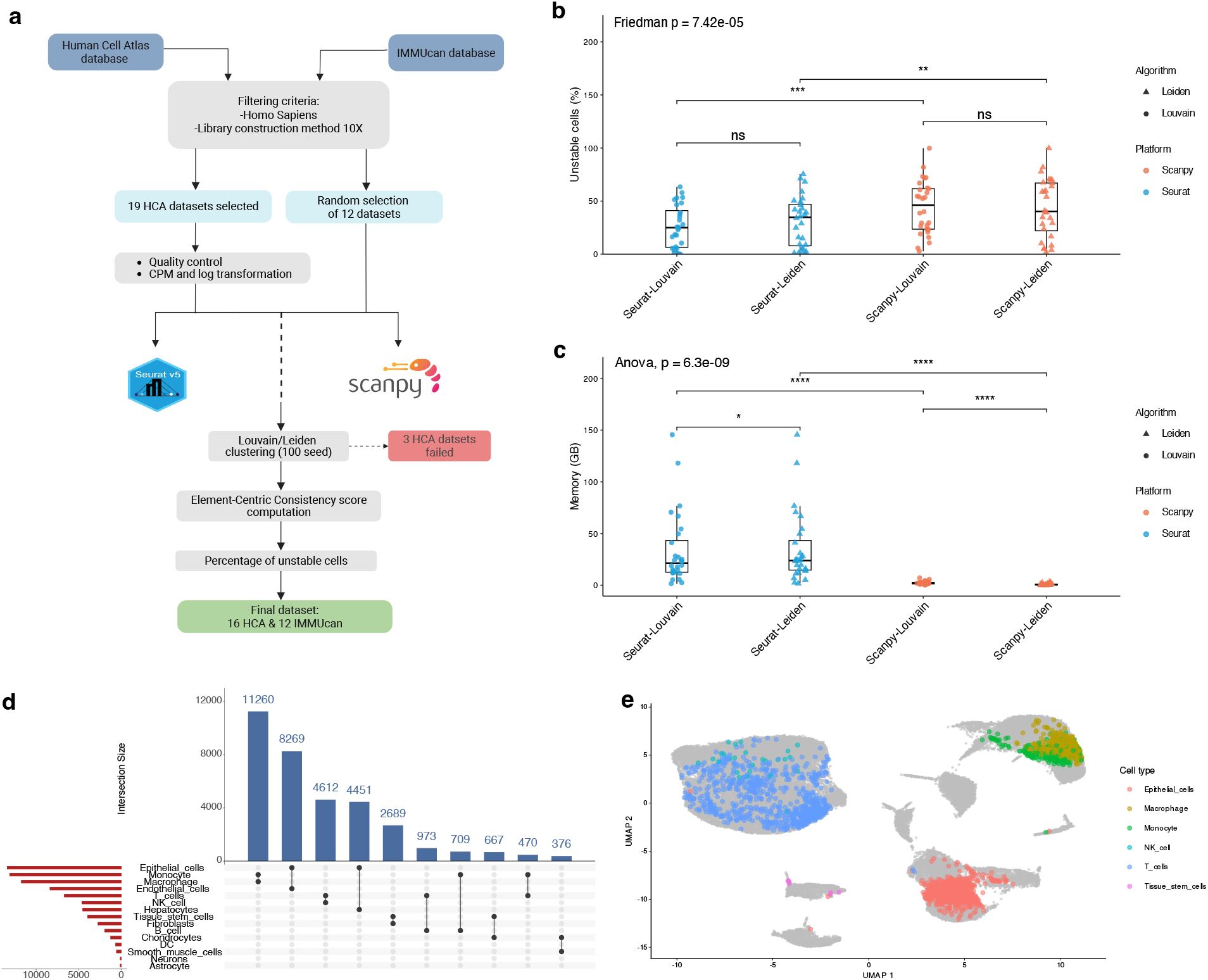
**a**, Overview of dataset selection and workflow for instability estimation. **b**, Percentage of unstable cells across platform–algorithm combinations; each point represents one dataset. Pairwise comparisons were performed using a paired Wilcoxon signed-rank test (* p<0.05; ** p<0.01; *** p<0.001) with BH correction, and overall differences were assessed by Friedman. **c**, Memory usage (GB) for each platform–algorithm combination. Pairwise comparisons were performed using a paired t-test (* p<0.05; ** p<0.01; *** p<0.001) with BH correction, and overall differences were assessed by ANOVA. **d**, Upset plot showing the co-occurrence of cell labels assigned to individual cells across IMMUcan datasets. Bars indicate intersection sizes and connected dots represent specific combinations of labels co-occurring. **e**, Example UMAP of the dataset NSCLC_UNB_10X_EMTAB6149 with unstable cells colored by cell type.

Seed variations produced substantial clustering instability across datasets and analytical settings. When Louvain and Leiden were considered jointly, Seurat generated more stable partitions than Scanpy, with a lower fraction of unstable cells (26.78%, interquartile range (IQR) 39.83) compared with Scanpy (40.46%, IQR 41.09; p-adj = 0.00000237; Fig. 1b, Supplementary Fig. 2, Supplementary Table 2). The most stable configuration was Seurat-Louvain, with a median of 25.06% (IQR 34.57) unstable cells, followed by Seurat-Leiden (34.75%, IQR 39.04), Scanpy-Leiden (40.26%, IQR 44.99), and Scanpy-Louvain (46.22%, IQR 38.18; Supplementary Table 3). Together, these results indicate that seed choice alone is sufficient to alter the assignment of a substantial fraction of cells, with instability ranging from a median of 25.06% in Seurat-Louvain to 46.22% in Scanpy-Louvain. This effect was not uniform across implementations, indicating that seed sensitivity is shaped not only by the clustering algorithm, but also by the analytical framework in which it is applied.

### Platform selection contributes to instability

We next asked whether seed-induced instability was driven primarily by the framework or by the choice of community detection algorithm. To separate these effects, we compared platform-level differences after averaging Louvain and Leiden within each dataset, and algorithm-level differences after averaging Seurat and Scanpy. The impact of the choice between Leiden and Louvain impacts only 5.96% of the unstable cells, while the impact of the platform (Seurat/Scanpy) affected 21.16%. This suggests that implementation framework contributes more strongly to seed-induced clustering instability than the choice between the clustering algorithm.

The observed increase of stability in Seurat came at a substantial computational cost. Across datasets, Seurat required markedly more memory than Scanpy for both Louvain and Leiden clustering (Fig. 1c; Supplementary Tables 4 and 5). After averaging Louvain and Leiden within platform, Seurat required 19 times more memory than Scanpy (p-adj = 9.55e-10), with a median memory usage of 23.44 GB (IQR 28.57) compared with 1.21 GB (IQR 1.8).

As shown in Supplementary Fig. 3 (Supplementary Table 6), memory usage increased linearly with the number of cells for both platforms, but the increase was markedly steeper in Seurat than in Scanpy. Across Louvain and Leiden, the average slope was ~1.00 GB per 1,000 cells for Seurat compared with ~0.035 GB per 1,000 cells for Scanpy, corresponding to an approximately 28-fold steeper memory scaling in Seurat. Both Louvain and Leiden showed the same overall pattern, for Louvain, Seurat required approximately 11-fold more memory than Scanpy; for Leiden, the increase was approximately 33-fold.

Although memory usage differed significantly between Louvain and Leiden within both Seurat and Scanpy, the largest differences were observed between frameworks. In particular, Seurat consistently required substantially more memory than Scanpy for both algorithms, indicating that platform-level implementation had a stronger impact on memory requirements than the choice between Louvain and Leiden. These findings reveal a trade-off: Seurat offers higher clustering stability at the cost of memory, highlight the importance of considering platform efficiency when working with large datasets, while Scanpy is more scalable but more sensitive to seed-dependent variation.

### Seed variation affects cluster annotation

To assess the biological impact of seed-dependent clustering instability, we performed automated annotation across 100 Seurat-Louvain clustering runs using the IMMUcan datasets. We showed that annotation instability was not uniformly distributed across cell types (Fig. 1d), inconsistent labels were enriched among biologically related or transcriptionally similar populations (Supplementary Table 7). The top 3 inconsistent label pairs were as follows: macrophage/monocyte-related switches represented 33.68% of inconsistent annotations, endothelial/epithelial populations the 29.51%, and T/NK cells the 15.32%.

A limited subset of cell-type labels accounted for the majority of annotation inconsistencies, suggesting that clustering variability preferentially affects populations with ambiguous transcriptional boundaries. This “fuzzy” behaviour can be further visualized in the example UMAP (Fig. 1e and Extended Data), where unstable cells tended to aggregate in defined regions of the embedding rather than being randomly distributed across the dataset. Overall, these results highlighted that seed variation in clustering is not only a technical source of clustering variability, but can also propagate to biologically meaningful downstream interpretations and affect the interpretation single-cell annotation results.

### SABER mitigates seed-induced instability through stability-based reassignment

To mitigate seed-induced clustering variability we developed StAbility-BasEd-Reassignment (SABER). The tool aims to achieve Seurat-like stability, despite its high memory demands, while retaining the low-memory advantages of the Scanpy framework. SABER identifies unstable cells through repeated clustering and reassigns them to stable clusters using cosine similarity. Stable clusters are defined from cells that are consistently assigned across repeated seed initialisations, allowing unstable cells to be reassigned on the basis of transcriptional similarity to robust cluster (Fig. 2a).

**Figure 2.**
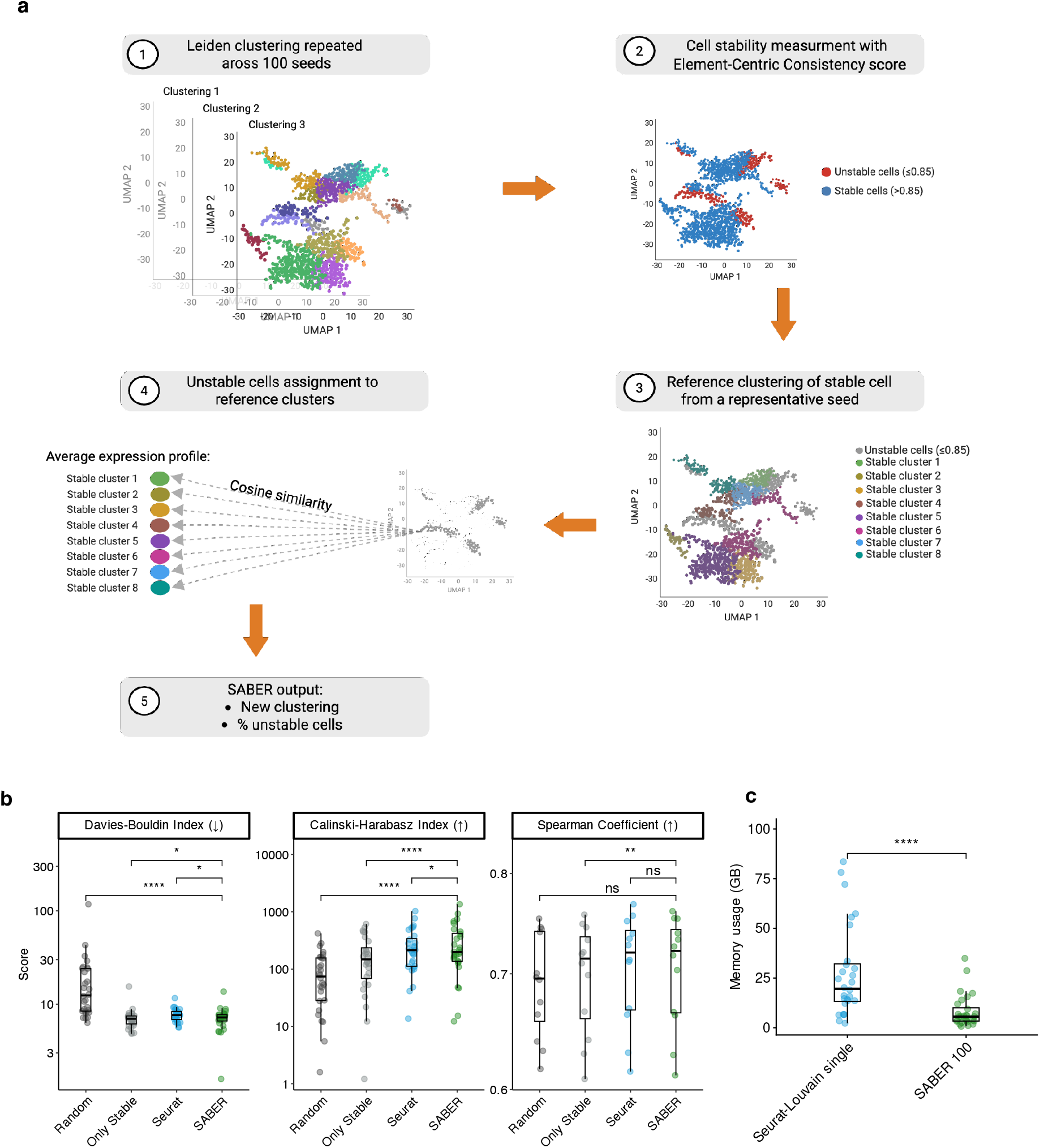
**a**, Overview of the SABER workflow. **b**, Comparison of SABER clustering with clusters obtained after random assignment of unstable cells to stable clusters, clusters with only stable cells, and cluster obtained with Seurat-Louvain. Statistical significance was assessed with paired Wilcoxon signed-rank test (* p<0.05; ** p<0.01; *** p<0.001) with BH correction. **C**, Distribution of memory usage (GB) of SABER and Seurat-Louvain single run. Significance was assessed by paired Wilcoxon signed-rank test) with BH correction.

We evaluated SABER performance against three alternative strategies: i) Random reassignment of unstable cells while preserving stable clusters sizes, ii) Exclusion of unstable cells followed by analysis of only stable cells, and iii) Seurat-Louvain clustering with a single arbitrary seed. These comparisons were chosen to capture complementary baselines: i) Random reassignment served as a negative control, testing whether SABER improved over chance-level allocation of unstable cells. ii) The stable-cell-only strategy represented a conservative alternative that removes unstable observations, but at the cost of reducing dataset size and statistical power. iii) Seurat-Louvain was used as a positive control because it represented the most stable configuration in our benchmark.

The three strategies were evaluated using the Davies-Bouldin (DB) index and the Calinski-Harabasz (CH) index, which measure cluster compactness and separation and, a weighted Spearman correlation for annotation concordance, computed only for the biologically homogeneous IMMUcan datasets, which allowed the use of a common reference for annotation.

SABER performed better than random reassignment, indicating that its cosine similarity-based reassignment captures meaningful structure rather than simply redistributing unstable cells by chance (paired Wilcoxon tests: DB, p-adj = 0.0000000223; CH, p-adj = 0.0000000223; weighted Spearman, p-adj = 0.064). SABER also outperformed the stable-cell-only strategy while retaining all cells, showing significant improvements in Davies-Bouldin index, Calinski-Harabasz score and weighted Spearman correlation (paired Wilcoxon tests: DB, p-adj = 0.016; CH, p-adj = 0.0000000336; weighted Spearman, p-adj = 0.007). These results indicate that stability-based reassignment is preferable to excluding unstable cells.

We then focused on the main benchmark against Seurat-Louvain, the most stable platform-algorithm combination identified in the seed variation analysis. Compared with a single Seurat-Louvain run, SABER achieved a lower Davies-Bouldin index, indicating improved cluster compactness and separation, with a median of 7.20 (IQR 1.12) compared with 7.63 (IQR 1.51) for Seurat-Louvain (p = 0.037). SABER showed a slightly lower Calinski-Harabasz score than Seurat-Louvain, with medians of 198.87 (IQR 286.14) and 213.93 (IQR 228.14), respectively (p = 0.012). However, annotation concordance was preserved: weighted Spearman correlation was comparable between SABER and Seurat-Louvain, with both methods reaching a median value of 0.72 (SABER IQR 0.08; Seurat-Louvain IQR 0.07; p = 0.176; Supplementary Table 8 and 9). Together, these results suggest that SABER improves cluster compactness while maintaining biological annotation consistency relative to the strongest stability baseline.

To further compare SABER with the variability of Seurat-Louvain across seed initializations, we computed Davies–Bouldin and Calinski-Harabasz scores for each of the 100 Seurat– Louvain runs and compared their dataset-level distributions with the corresponding SABER partition. Using the median value across Seurat-Louvain initializations as reference, SABER improved the Calinski-Harabasz score in 19 of 28 datasets (67.9%) and the Davies–Bouldin index in 18 of 28 datasets (64.3%) (Supplementary Table 10). This indicates that SABER is not only noninferior to a single Seurat-Louvain run, but often performs better than the typical Seurat-Louvain partition obtained across seed initializations.

Then, we quantified the agreement between SABER and 100 Seurat–Louvain partitions using Normalized Mutual Information (NMI) and Adjusted Rand Index (ARI). Across datasets, SABER showed a median NMI of 0.61 (IQR 0.17) and a median ARI of 0.37 (IQR 0.20) when compared with the 100 Seurat-Louvain initializations (Supplementary Table 11). The higher NMI compared with ARI suggests that SABER preserves part of the global information shared with the original Seurat-Louvain partitions, while producing local changes assignments that reduce pairwise agreement. This behaviour is expected because SABER is not designed to reproduce any seed specific partition, but rather to retain the stable clustering structure shared across seed initializations and modify the final partition through reassignment of unstable cells.

Overall, these results show that SABER provides a robust and biologically consistent strategy to reduce seed-induced clustering variability. By reassigning unstable cells rather than discarding them, SABER preserves the full dataset while improving clustering quality.

### SABER improves scalability while preserving clustering quality

Because one of the main goals of SABER was to provide a scalable alternative to memory-intensive Seurat workflows, we compared the computational requirements of the full SABER pipeline with a single Seurat-Louvain run. SABER required significantly less memory, than a single run of Seurat-Louvain (Fig. 2c), despite incorporating 100 seed iterations. Median memory usage was 5.58 GB (IQR 6.69) for SABER compared with 19.59 GB (IQR 18.91) for Seurat–Louvain, corresponding to a 3.5-fold reduction in median memory usage (p-value = 0.00000126; Supplementary Table 12). This result indicates that SABER combines two advantages: it explicitly quantifies and mitigates seed-induced clustering instability, while maintaining the computational scalability of the Scanpy framework. In this way, SABER provides an optimal compromise between the stability observed in Seurat-based clustering and the lower memory requirements of Scanpy.

## Discussion

Clustering is a central step in scRNA-seq analysis, yet its sensitivity to stochastic initialization is rarely evaluated in practice. By varying seeds across platforms (Seurat and Scanpy), algorithms (Louvain and Leiden), and datasets, we show that seed choice alone can alter cell assignments. These results demonstrate that clustering outcomes commonly treated as deterministic are dependent on seed, with consequences for reproducibility and biological interpretation.

Across 28 datasets, Seurat exhibited more stable clustering results than Scanpy for both Louvain and Leiden algorithms. However, this increased stability comes with a higher computational cost, approximately 11-fold higher for Louvain and 33-fold higher for Leiden. In contrast, Scanpy exhibited superior scalability and lower memory requirements, but greater susceptibility to seed variability. This trade-off highlights a fundamental tension between computational efficiency and clustering robustness in current single-cell analysis frameworks.

Clustering instability was not uniformly distributed across datasets or cell populations. Our analysis of annotation instability revealed that specific cell populations, often transcriptionally similar, are particularly sensitive. For example, macrophages and monocytes, endothelial and epithelial cells, and subsets of T and NK cells accounted for the majority of inconsistent annotations. These findings suggest that stochastic effects are amplified when biological boundaries are inherently ambiguous. As cluster annotations are often propagated to downstream analyses, including differential expression and trajectory inference, such instability has the potential to introduce systematic biases.

To address these challenges, we developed StAbility-BasEd-Reassignment (SABER), a post hoc framework that leverages multiple clustering iterations to identify unstable cells and reallocates them to stable clusters based on expression similarity. SABER effectively balances cluster stability and biological relevance, retaining all cells while improving clustering quality relative to random assignment and standard Seurat-Louvain clustering. Although SABER relies on multiple clustering iterations, it is substantially less memory intensive than a single Seurat-Louvain run and achieves comparable stability to Seurat, supporting its use in large scale or resource limited settings.

In summary, our study demonstrates that random seed variation is a source of variability in single-cell clustering analyses. By providing both a framework to assess clustering instability and a scalable method to mitigate its effects, SABER offers a practical solution to enhance robustness and reproducibility in scRNA-seq workflows. These findings underscore the importance of explicitly accounting for stochasticity in clustering and suggest that stability-aware approaches should become a standard component of single-cell data analysis.

## Methods

### Datasets

To evaluate the impact of seed variation on clustering stability, we collected publicly available single-cell RNA-seq datasets. We searched the Human Cell Atlas (HCA) portal (https://data.humancellatlas.org/), filtering for *Homo sapiens*, and 10X library construction protocols. We retained a total of 19 scRNA-seq datasets. Additionally, 12 randomly selected datasets were downloaded from the IMMUcan database (https://immucanscdb.vital-it.ch/), these datasets were already filtered and normalized. Detailed dataset descriptions and download links are available in Supplementary Table 1.

### Preprocessing and clustering

Single-cell RNA-seq datasets from HCA were subjected to quality control using Scanpy. For each sample, standard QC metrics were computed, including the number of detected genes and the percentage of mitochondrial reads. Filtering thresholds were defined using a median absolute deviation (MAD) approach (±3 MAD for the number of detected genes and upper 3 MAD for mitochondrial content). Cells failing these criteria were excluded. Filtered datasets were saved individually and subsequently concatenated using the sample identifier as batch annotation. A scRNA-seq analysis pipeline was developed, comprising the following steps: i) Count Per Million normalization, ii) logarithmic transformation of the counts, iii) PCA, iv) integration, v) clustering using both the Louvain and Leiden algorithms. We performed integration using Harmony applied to the principal component space. The batch variable was defined as the sample (or Sample) metadata field when available. In both Scanpy and Seurat pipelines, Harmony was run after PCA computation and the corrected embeddings were used for all downstream analyses. When no batch annotation was available, batch correction was not applied. The clustering process was repeated 100 times, with seed values ranging from 1 to 100. The analysis was conducted using Seurat (5.3.0) in R (4.4.3) and Scanpy (1.11.4) in Python (3.10.19), with a resolution of 0.6 to ensure robustness across platforms. We worked on a virtual machine with 32 CPU cores, 190 GB of RAM memory, 60 GB of swap memory.

For cross-platform consistency, we applied fixed parameter settings across all datasets, using the respective default implementations in Seurat and Scanpy. The normalization step was performed only for HCA datasets, while IMMUcan counts were already normalized; for both Seurat and Scanpy we used a scale factor of 1×10^6^ and a log-transformed with log(1+x). The first 2000 Highly variable genes (HVGs) were identified in both pipelines with their default methods, selection.method=“vst” option in Seurat and flavor=“seurat” in Scanpy. Prior to clustering, we performed PCA with default number of components of 50 in both platforms and then we computed nearest-neighbor graph with FindNeighbors (reduction = “pca”, dims = 1:20) in Seurat which construct a Shared Nearest-neighbor; while we found *k-*nearest neighbours with sc.pp.neighbors (n_pcs=20) in Scanpy. These choices allow systematic comparison across multiple datasets, though they may not represent the optimal parametrization for each individual dataset.

### Memory measurement

To evaluate computational requirements, we measured memory usage for each dataset in both platforms. In R/Seurat, benchmarking was performed using the peakRAM package (1.0.3), which reports memory (bytes) allocation during the execution of the pipeline.

Memory usage was quantified by capturing peak memory allocation (bytes) using tracemalloc (3.10.18). The collected statistics were saved for each dataset to allow systematic comparison of computational costs across tools and datasets. To quantify how memory usage scaled with dataset size, we fitted a linear regression model. Specifically, for each platform and clustering algorithm, memory consumption was modeled as a function of the number of cells in the dataset: *memory* ~ *number of cells*. The slope of the resulting regression line represents the rate of increase in memory usage per additional cell, providing a measure of how efficiently each platform scales with dataset size.

For evaluating SABER scalability, we measured memory using a bash script. Memory usage was monitored by summing the resident set size (RSS) of all child processes spawned by the main pipeline, and the maximum observed value was recorded as peak memory usage.

### Identification and measurement of the cells unstable to seed variation

To quantify cell instability across runs, we employed Element-Centric Consistency (ECC) score, implemented in the ClustAssess R package (1.1.0)^17^. ECC was selected because it enables to assess for each cell its clustering stability, which is critical for identifying individual cells that are sensitive to stochastic variation. The ECC score, which ranges from 0 to 1, is a measure of the consistency with which cells are grouped together across different clustering iterations. The ECC is measured for each element (cell), starting with the computation of a personalized probability distribution (PPR), which reflects the probability of that element belonging to each cluster in a particular clustering. Next, for each pair of clustering, the L1 norm distance is computed between their PPRs. This L1 distance measures the sum of the absolute differences between the corresponding PPR values for each element. After calculating the L1 distances for every unique pair of clustering, these L1 scores are summed for each element and then divided by the total number of comparisons. This averaging step produces a final consistency score for each element, representing the mean stability of its cluster assignments across all clustering pairs. The normalization ensures that the final score lies between 0 and 1.

### Seed variation effect on cluster annotation

To assess the effect of seed variation on cluster annotation, we evaluated the consistency of SingleR^18^ (2.8.0) package annotations across multiple clustering obtained varying the seed from 1 to 100 in the 12 IMMUcan datasets. For each dataset, we performed 100 clustering iterations using Seurat-Louvain. The reference used was the Human Primary Cell Atlas Data from the celldex package (1.16.0). For each clustering iteration, SingleR was applied at the cluster level, and the resulting labels were then propagated back to individual cells. For each cell, we collected the set of unique labels assigned across the 100 runs. Cells receiving more than one distinct annotation label were classified as mislabeled cells. To further characterize annotation variability, we analyzed the frequency and overlap of cell labels assigned to cells across all datasets.

### ECC threshold determination

To identify cell instability, we combined for each cell the annotation frequencies with the ECC score obtained with Seurat-Louvain. Cells that changed annotation in more than 5% of clustering runs were considered unstable annotated. For each dataset, we calculated the 95th percentile of ECC values among these cells, generating a dataset-specific threshold. To obtain a single, global threshold applicable across all datasets, we calculated the median of these dataset-specific thresholds, obtaining a value of 0.85. Cells with ECC scores below or equal this threshold were classified as ‘unstable’. This approach linked ECC thresholds to annotation instability rather than an arbitrary cutoff.

### StAbility-BasEd-Reassignment (SABER) implementation

To identify and correct seed variability in clustering results, we developed SABER (StAbility-BasEd-Reassignment) method to reassign the unstable cells previously identified.

### Reassignment of unstable cells

The workflow started with clustering performed 100 times with different seeds using the Leiden algorithm in Scanpy. For each cell, the ECC was computed across all 100 clusterings. Cells with an ECC > 0.85 were considered stable, while those below this threshold were defined as unstable. Because different seeds could yield varying numbers of clusters, we identified the most frequently observed cluster count across the 100 runs, then selected the seeds that produced this count and randomly selected one for re-clustering only the stable cells. The stable cells were then clustered again using the Leiden algorithm with a resolution of 0.6, and the resulting clusters were defined as stable clusters. To re-assign unstable cells, each cell’s expression profile was compared to the mean expression profile of the stable clusters, and the cell was assigned to the cluster with highest cosine similarity.

### Dataset for SABER evaluation

All the datasets were used to evaluate the effectiveness of the SABER method using Davies-Bouldin index and the Calinski-Harabasz index. Weighted Spearman correlation coefficients were computed exclusively for 12 datasets from IMMUcan. We selected these immunological datasets to enable automated cluster annotation using the SingleR (2.8.0) package, with the Human Primary Cell Atlas Data from the celldex package (1.16.0) serving as reference.

### Clustering quality assessment

The quality of the re-allocation was assessed using two clustering validation metrics, that provide complementary internal assessments of clustering quality without requiring external labels: the Davies-Bouldin index and the Calinski-Harabasz index from the sklearn (1.7.1) package.

The Davies-Bouldin index (DB) assesses the average similarity between each cluster and the most similar one among the others. It takes into account both intra-cluster compactness and inter-cluster separation and is defined as:

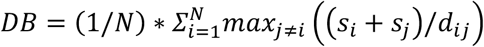

where *N* is the number of clusters, *s*_*i*_ the average intra-cluster distance for cluster *i* and *d*_i*j*_the distance between centroids of clusters *i* and *j*. The DB index ranges from 0 to infinite, with lower values indicating more compact and better-separated clusters.

The Calinski-Harabasz index (CH), on the other hand, measures the ratio of inter-cluster dispersion to intra-cluster dispersion, and is defined as:

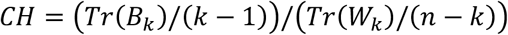

where *k* is the number of clusters, *n* the total number of points, *Tr*(*B*_*k*_) the trace of the between-cluster scatter matrix and *Tr*(*W*_*k*_) the trace of the within-cluster scatter matrix. The

CH index is unbounded, with higher values corresponding to better-defined clusters, where data points are more spread out between clusters than they are within clusters.

### Clusters evaluation using Spearman coefficient

For each annotated cluster and cell type present in the reference, Spearman correlation coefficients were computed by SingleR to assess the similarity:

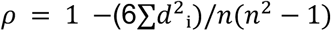

where *d*^2^_i_ is the difference between the ranks of each pair of observations and *n* is the number of points. For each cluster, the highest Spearman coefficient across all the reference cell types was retained. To provide an overall measure of annotation quality for each dataset, a weighted mean of the Spearman coefficients was calculated:

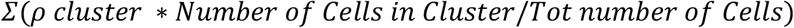

This metrics summarizes of how well the annotated clusters correspond to the reference profiles while accounting for differences in cluster size.

### Metric comparison and benchmarking

Clustering performance was evaluated using the DB index, CH index, and the weighted Spearman correlation coefficient. These metrics were used to assess the effectiveness of the SABER method and to compare it with three alternative clustering strategies: i) random reassignment of unstable cells while preserving stable cluster sizes, ii) clustering based exclusively on stable cells and iii) Seurat-Louvain clustering using an arbitrary seed.

To further evaluate clustering quality at the dataset level, DB and CH indices were computed for all the Seurat-Louvain clustering generated by varying the seed from 1 to 100. Each of these scores was then compared with the corresponding value obtained using SABER. Agreement between clustering results produced by SABER and those obtained from Seurat-Louvain across different seeds was quantified using the Normalized Mutual Information and Adjusted Rand Index. For each dataset, the SABER clustering was used as the reference and compared against each of the 100 Seurat-Louvain clusterings.

The Normalized Mutual Information (NMI) is a normalized version of the Mutual Information (MI) score that quantifies the amount of information shared between two clustering. NMI ranges between 0 and 1, where 0 indicates no mutual information and 1 indicates identical clustering. NMI was computed using the normalized_mutual_info_score function from sklearn. The NMI formula is:

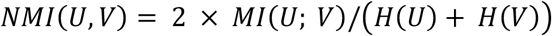

where MI(U; V) is the mutual information between U and V, and H(U) and H(V) are the entropies of U and V, respectively.

The Adjusted Rand Index (ARI) measures the similarity between two clustering by evaluating the agreement of all pairs of cells assigned either to the same or different clusters. ARI values range from −1 to 1, where 1 indicates perfect concordance. ARI was computed using the adjusted_rand_score from sklearn. The ARI formula is:

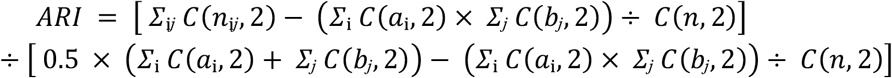

where n_ij_ is number of elements in common between cluster i in U and cluster j in V, a_i_ = Σ_j_ n_ij_, b_j_ = Σ_i_ n_ij_, n is the total number of elements and C(x, 2) = combinations of x elements taken 2 at a time = x·(x−1)/2.

### Statistical analysis

The choice of statistical tests was guided by the assessment of data normality using the Shapiro-Wilk test. Parametric tests, including the t-test and ANOVA, were applied to variables satisfying normality assumptions. When the normality assumption was not met, the Wilcoxon rank-sum test was used for pairwise comparisons and the Friedman test for multiple-group comparisons. The tests were corrected with Benjamini-Hochberg (BH) procedure.

## Data availability

The datasets used in this study were downloaded from https://explore.data.humancellatlas.org/projects and https://immucanscdb.vital-it.ch/. All the links to access the datasets are available in the Supplementary Table 1.

## Code availability

All code used for the analyses is deposited in a GitHub repository (https://github.com/mazzalab-bioinfo/SABER-seed-variability), providing both the scripts required to reproduce the study and detailed instructions for running SABER using Docker.

## Acknowledgements

N.Z. is a PhD student within the European School of Molecular Medicine (SEMM).

E.B. was supported by Fondazione Umberto Veronesi.

## Contributions

NZ, GT, EB and LM contributed to conceptualization. NZ, GT and EB contributed to methodology. NZ developed the bioinformatic pipelines and performed the formal analysis, investigation, data curation and visualization, and wrote the original draft of the manuscript. GT, EB and LM supervised the study and reviewed and edited the manuscript. LM acquired funding. All authors reviewed and approved the final manuscript.

## Funding

-Italian Ministry of Health with Ricerca Corrente and 5 × 1000 funds

-European Union – Next Generation EU – PNRR M6C2 – Investimento 2.1 Valorizzazione e potenziamento della ricerca biomedica del SSN – Project Code: PNRR-MAD-2022

-Ministry of Health “Giovani Ricercatori” grant GR-2019-12369910

-AIRC MFAG n 25791.

-This research was supported by the Italian Ministry of Health, under the “PSC Salute, Traiettoria 4” program, specifically the “CAL.HUB.RIA” project (code T4-AN-09).

