## Supplementary Fig for "Seed variation impacts clustering stability in Single-Cell RNA-Seq and can be mitigated by StAbility-BasEd-Reassignment (SABER)"

**Supplementary Figures**

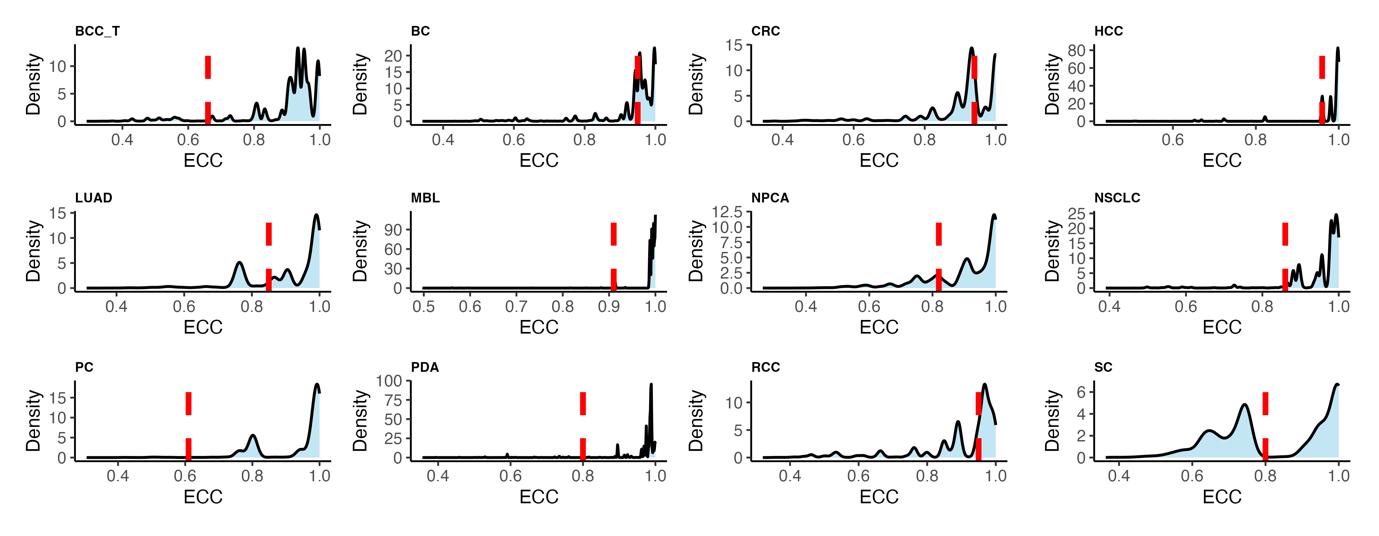
**Supplementary Figure 1 | ECC threshold in IMMUcan datasets**

Density distribution of ECC scores computed for individual cells across 100 Seurat-Louvain clustering iterations in the IMMUcan datasets. The red dashed line indicates the 95th percentile of ECC distribution among unstable cells (those changing annotation in more than 5% of clustering runs). The median of these values was used to define the final threshold used to define unstable cells.

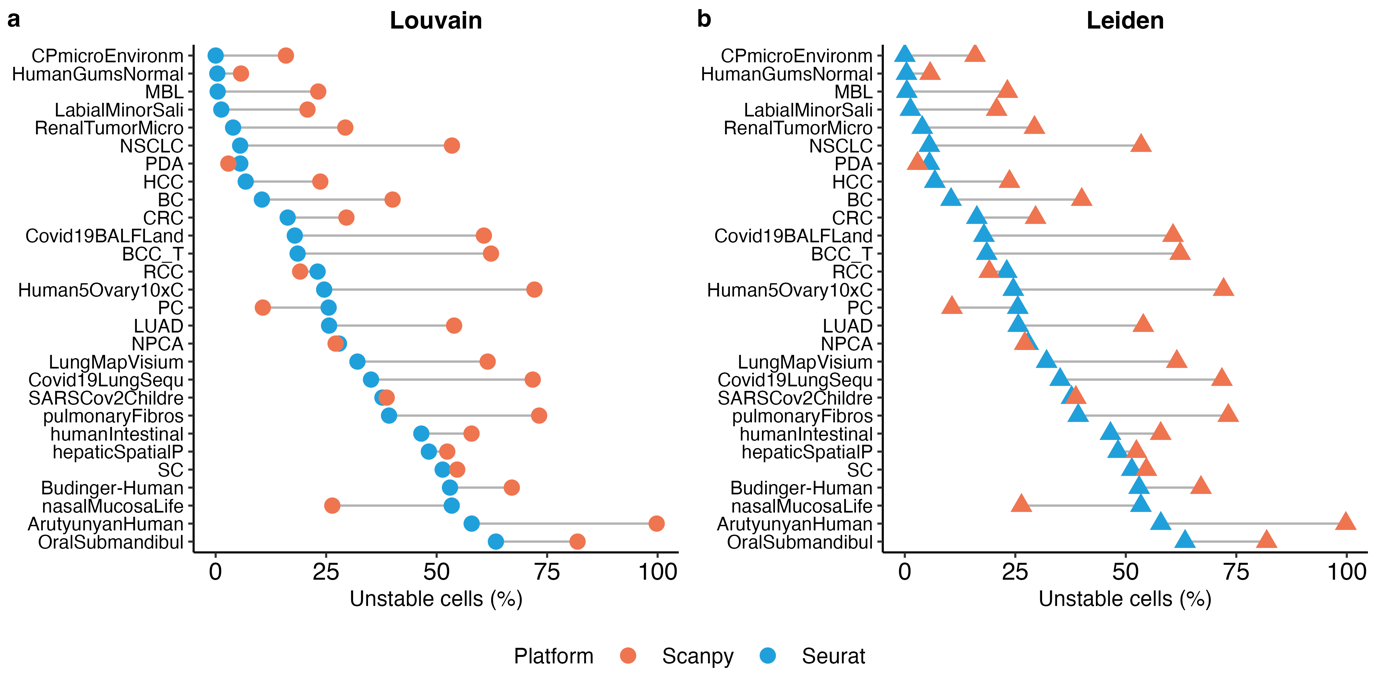

**Supplementary Figure 2 | Comparative performance of Seurat and Scanpy.
a, b,** Percentage of unstable cells with ECC values ≤ 0.85 identified by Seurat and Scanpy using the Louvain (a) and Leiden (b) algorithms across100 seed iterations.

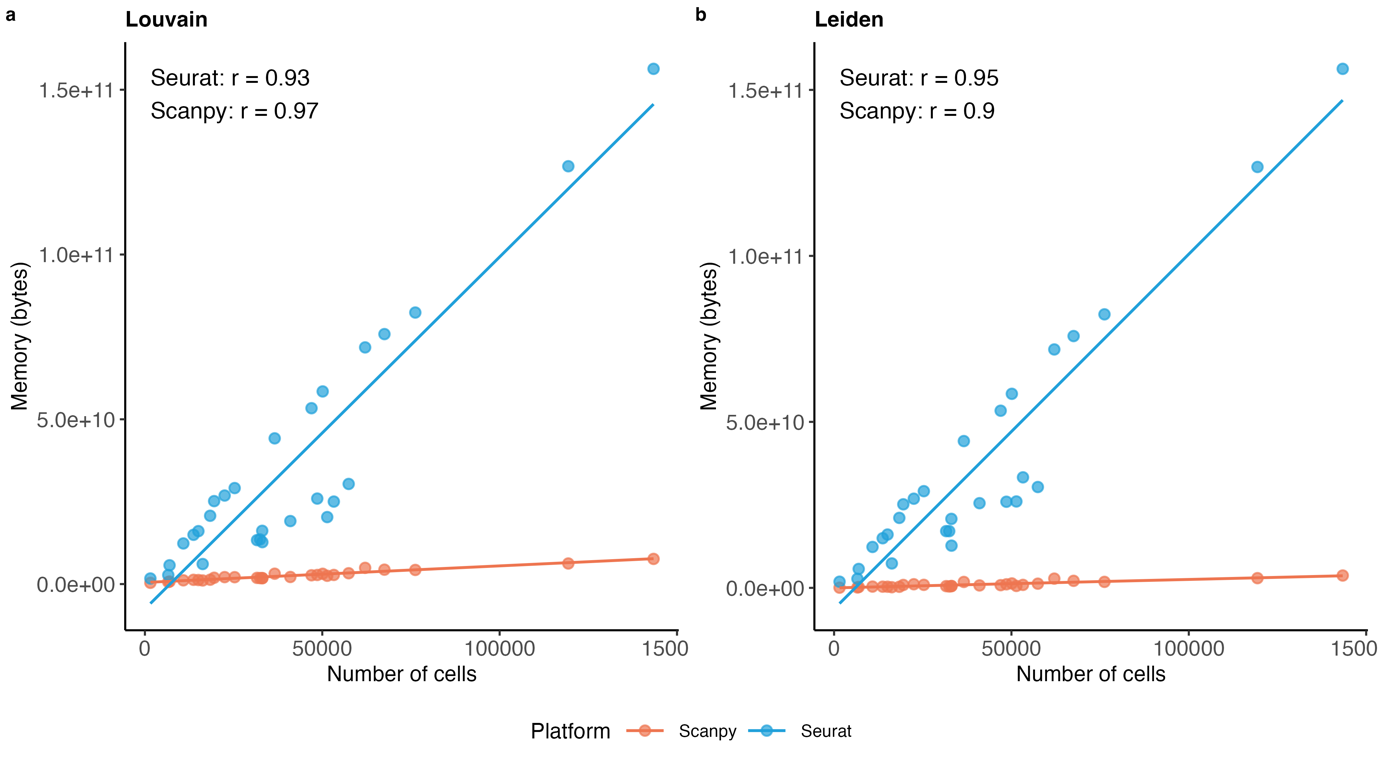

**Supplementary Figure 3 | Correlation between memory usage and number of cells**

**a, b**, Scatter plots show memory consumption (bytes) versus number of cells for Seurat and Scanpy using the Louvain (a) and Leiden (b) algorithms. Solid lines indicate linear regression fits. Pearson correlation coefficients (r) between memory usage and dataset size are reported within each panel.
